# Distinct mechanisms underlie value-driven modulation of visual cortex by previously rewarded visual and auditory stimuli

**DOI:** 10.1101/2023.01.25.525484

**Authors:** Jesssica Emily Antono, Shilpa Dang, Ryszard Auksztulewicz, Arezoo Pooresmaeili

**Affiliations:** Perception and Cognition Lab, European Neuroscience Institute Goettingen- A Joint Initiative of the University Medical Center Goettingen and the Max-Planck-Society, Germany, Grisebachstrasse 5, 37077 Goettingen, Germany; School of Artificial Intelligence and Data Science, Indian Institute of Technology Jodhpur, Hauz Khas, New Delhi-110016, India; Center for Cognitive Neuroscience Berlin, Free University Berlin, Habelschwerdter Allee 45, 14195 Berlin, Germany

**Keywords:** reward, value, visual perception, sensory modality, fMRI

## Abstract

Past reward associations may be signaled by stimuli from different sensory modalities, however it remains unclear how different types of reward-associated stimuli modulate perception. In this human fMRI study, we employed a paradigm involving a visual discrimination task, where a visual target was simultaneously presented with either an intra-(visual) or a cross-modal (auditory) cue that was previously associated with rewards. We hypothesized that depending on the sensory modality of the cues distinct neural pathways underlie the value-driven modulation of visual areas. Two steps of analyses were conducted: first, using a multivariate approach, we confirmed that previously reward-associated cues enhanced the target representation in the early visual areas. Then, using effective connectivity analysis, we tested three possible patterns of communication across the brain regions that could underlie the modulation of visual cortex: a direct pathway from the frontal valuation areas to the visual areas, a mediated pathway through the attention-related areas, and a mediated pathway that additionally involved distinct sensory association areas for auditory and visual rewards. We found evidence for the third model and demonstrate that reward-related information is communicated across the valuation and attention-related brain regions such as the intraparietal sulcus across for both visual and auditory cues. Additionally, the long-range communication of reward information also involved the superior temporal areas in case of auditory reward-associated stimuli. These results suggest that in the presence of previously rewarded stimuli from different sensory modalities, a combination of domain-general and domain-specific mechanisms are recruited across the brain to adjust visual processing.

## Introduction

Rewards modulate information processing in the brain at multiple stages, from decision making where an organism’s behavior is optimized to maximize reward outcomes (O’Doherty et al., 2001, 2007), to perception where the representations of sensory stimuli are altered depending on their current or past associations with rewards (Cicmil et al., 2015; Hickey et al., 2010; Rangel et al., 2008; Serences, 2008; Stanisor et al., 2013; Arsenault et al., 2013). Previous literature has demonstrated that a network encompassing the ventral striatum and prefrontal cortex plays a crucial role in learning and representation of reward value, thereby informing the subsequent decision-making stages about the best course of action to choose (Schultz, 2000; Rangel et al., 2008). On the other hand, a more recent line of research has provided evidence for a value-driven modulation of neuronal responses in almost all primary sensory areas (Rutkowski and Weinberger, 2005; Shuler and Bear, 2006; Pleger et al., 2008; Weil et al., 2010; Goltstein et al., 2013; Stanisor et al., 2013), a mechanism through which stimuli associated with higher rewards or better realization of the goals of the task are prioritized for perceptual processing. Despite the wealth of knowledge regarding the neuronal underpinnings of valuation in the brain and the emerging evidence for the value-driven alteration of perception, it is unclear how these processes interact.

Unravelling the mode of interaction between valuation and perception is a crucial step towards understanding how information processing in the brain is adapted to the rich and dynamic characteristics of the naturalistic environments. In such settings, objects have multiple features; from the same or different sensory modality; which may have different associations with rewards, and these associations may change over time. Therefore, to form a robust representation of reward value despite the multitude of stimulus features in the environment, the valuation network should constantly receive information from sensory areas (Komura et al., 2001; Reig and Silberberg, 2014). On the other hand, sensory areas should be efficiently re-regulated as reward associations of stimuli and task requirements undergo changes so that in each instance the stimuli that lead to better outcomes gain advantaged processing (Haber, 2011; Pessoa and Engelmann, 2010).

Different models have been put forward to explain the communication of information across the brain’s valuation network and the sensory areas. Pessoa & Engelmann (2010) for instance, proposed that reward signals are embedded into perceptual processing through either direct or indirect inputs from the valuation network to sensory areas. Direct inputs rely on a connectivity between the valuation network and sensory areas, whereas indirect inputs are likely to be first broadcasted to the frontoparietal attentional network (Corbetta and Shulman, 2002; Pessoa, 2009) and then be fed back to the sensory areas. Additionally, recent studies have identified other sensory association areas which may be involved in routing information between the valuation and perception networks. For instance, Pooresmaeili et al., (2014) found an increase of neural responses in the superior temporal cortex, known to be involved in multisensory processing (Calvert et al., 2000, 2001; Stein and Stanford, 2008), when auditory stimuli had been associated with higher rewards and modulated visual perception cross-modally. This finding suggested that areas involved in combining information across different features of a multisensory object may additionally integrate reward signals into the perceptual processing (Cheng et al., 2020). This proposal is also in line with the findings from another study (Anderson, 2017) showing that lateral occipital complex (LOC), an area that is involved in representation of perceived objects (Kourtzi and Kanwisher, 2001) and integration of local features to global shapes (Grill-Spector, 2003) especially when attention is biased to visual object features (Martin et al., 2018), plays a role in the value-driven changes in attentional control. Yet another possibility is that a history of privileged processing and preferred selection confers high reward stimuli a long-lasting processing gain already at the level of encoding of information at the early visual areas (Kim and Anderson, 2019), and hence value-driven modulation of perception occurs without the need for constant communication of information across the valuation and perception systems.

All mechanism outlined above have found support in the literature. For instance, direct inputs from the valuation network is plausible because previous studies have shown that lateral OFC and striatum have bilateral connections with the primary visual cortices (Barbas, 1993; Carmichael and Price, 1995; Kveraga et al., 2007; Khibnik et al., 2014). However, these connections may first be relayed to other areas as direct dopaminergic inputs to early visual areas such as area V1 are scarce (Oades and Halliday, 1987; Jacob and Nienborg, 2018) therefore supporting the proposal of a mediation through the sensory association (Macedo-Lima and Remage-Healey, 2021) or attentional (Noudoost and Moore, 2011) areas. The important role of attentional areas in mediation of value-driven effects is also supported by a host of previous studies (Pessoa, 2015), demonstrating that rewards guides attention to be allocated to the most valuable items in the scene (Chelazzi et al., 2013), and attention in turn gates the effects of reward by determining whether or not rewarded stimuli are aligned with the goal of the task and should be boosted or supressed (Roelfsema and Van Ooyen, 2005; Roelfsema et al., 2010; Gong et al., 2017). Finally, an effect of reward locally arising at the level of sensory areas due to the reward history and its resultant long-lasting changes in sensory representations is supported by computational modelling (Wilmes and Clopath, 2019) and experimental approaches (Chubykin et al., 2013; Kim and Anderson, 2019) showing that during learning, the task-relevant neural representations that are predictive of rewards are locally boosted in area V1 (Poort et al., 2015).

The aim of the current study was to shed light on the underlying mechanisms of value-driven modulation of perception and the mode of interaction between the valuation and perception systems. Specifically, we sought to test which of the mechanisms mentioned above can best explain the value-driven modulation of visual perception across different types of reward-associated stimuli. Towards this aim, we used a behavioral paradigm similar to previous studies (Pooresmaeili et al., 2014; Vakhrushev et al., 2021; Antono et al., 2022) that featured either cross-modal (Pooresmaeili et al., 2014) or both cross- and intra-modal reward-associated stimuli (Vakhrushev et al., 2021; Antono et al., 2022). In this paradigm, auditory or visual stimuli were first associated with either high or low monetary reward during a reward associative learning phase (referred to as conditioning). During the test phase (post-conditioning), auditory and visual reward-associated stimuli (cross- and intra-modal, respectively) were presented at the same time as the target of a visual discrimination task but were irrelevant to the task at hand and did not predict the delivery of reward anymore. By having a comparison between intra- and cross-modal reward associated cues, we aimed to identify reward-related mechanisms that are shared or disparate across sensory modalities. Furthermore, in order to disentangle reward- and goal-related mechanisms, we associated rewards to the features of the stimulus that were not the target of the visual discrimination task. Concurrently as participants performed the behavioral task, we recorded the brain activity using fMRI.

We hypothesized that higher reward improves performance by enhancing the neural representation of the task target in the early visual areas. In our task, the visual discrimination had to be done on a target stimulus (i.e., a Gabor patch) while the reward-associated stimuli were presented simultaneously and at the same spatial location but were irrelevant to the task. Therefore, to examine the target-specific modulation of visual processing, we inspected how the accuracy of a multivariate pattern classifier for target’s tilt orientation in the early visual areas was influenced by the value of reward-associated stimuli. Furthermore, to identify which brain areas were engaged in encoding the associated reward value of stimuli, we used a second set of multivariate pattern classifiers that decoded stimulus value, either dependent or independent of specific sensory features, across the brain. Finally, we tested possible models of whether and how the long-range communication of reward information between the valuation and early visual areas occurs. Our results showed that overall higher reward enhanced the accuracy of target-specific representations in the early visual areas but this effect involved distinct modes of long-range neuronal interactions across the brain for cross-modal and intra-modal reward-associated stimuli.

## Materials and Methods

### Participants

Thirty-six healthy participants were recruited (14 females; mean age 25.6 ± 4.48 SD, 20-40 years old) using an online local database (http://www.probanden.eni-g.de/orsee/public/). All participants had normal or corrected-to-normal vision, were right-handed, and gave oral and written informed consent after all procedures was explained to them. 3 participants were excluded from all analyses since their performance in the reward conditioning task was below a pre-defined criterion (<80%) indicating that they could not localize the visual or auditory stimuli. One additional participant was excluded from the fMRI analysis since the data acquisition inside the scanner could not be completed (see the *Procedures*). Participants were paid 10€ per hour for their participation in 2 scanning sessions (each 2.5 hours), and in addition received a bonus up to 10€ depending on their performance. The study was approved by the local ethics committee of the “Universitätsmedizin Göttingen” (UMG), under the proposal number 15/7/15.

### Stimuli and apparatus

The target stimuli used for the main task in the pre- and post-conditioning were Gabor patches (a Gaussian-windowed sinusoidal grating with SD = 0.33°, a spatial frequency of 3 cycles per degree, subtending 2° diameter, displayed at 10° eccentricity to the left or right side of the fixation point), which were tilted clockwise or counter-clockwise relative to the horizontal meridian. In each trial, a semi-transparent ring (alpha 50%, 0.44° in diameter) was superimposed on the Gabor patch. The color of the rings (orange or blue for visual conditions, or grey for auditory and neutral conditions) was adjusted individually for each participant in such a way that they were perceptually isoluminant. For auditory cues, two pure tones (600 Hz or 1000 Hz) were presented at 90dB simultaneously with the Gabor patch and at the same spatial location (see the *Procedures*). To achieve the co-localization of the auditory tones and the visual stimuli, we convolved the tones with head-related transfer functions based on a recorded database (Algazi et al., 2001) so that they could be perceived at 10° distance to the left or right of the fixation point.

Throughout the experiment, visual stimuli were displayed on an MR-compatible projection screen using a calibrated ProPixx projector (VPixx Technologies, Saint-Bruno, QC, Canada) at a resolution of 1920×1080 pixels, and a refresh rate of 120 Hz. The screen was placed at the end of the scanner bore at a distance of 88 cm from the participants’ eyes. The full display size on the screen was 43 cm x 24 cm, i.e. the visible range from the central fixation spot was +/-13.6° horizontally and +/-7.7° vertically. The auditory tones were delivered through MR compatible earphones (Sensimetric S15, Sensimetrics Corporation, Gloucester, MA) with an ear tip (Comply™ Foam Canal Tips) to maintain acoustic seal and reduce environmental noise.

For tracking the gaze position an MRC eye-tracker system mounted on the mirror on top of the MR head coil was used (MRC HiSpeed, MRC Systems GmbH, Heidelberg, Germany). Before each of the two scanning sessions, the eye-tracking system was calibrated using a 9-point standard MRC calibration procedure.

### Procedures

The data collection was done over two scanning sessions (about 2.5 hours each). The first session consisted of a preparation phase (comprising a practice session for the visual orientation discrimination task: VDT, measurements of the sound localization, adjustment of colors’ luminance and determining the perceptual threshold for the VDT) and an experimental phase referred to as pre-conditioning with the simultaneous acquisition of fMRI data. Prior to the scanning, participants completed a sound localization task, where they had to indicate whether a sound was played from the left or right side using their index and middle finger on a keyboard, and were included in the study if their localization accuracies were >95%. Afterwards, participants adjusted the luminance of both visual cues using a flicker task inside the scanner. The tilt orientation of the Gabor patch during the orientation discrimination task was set to each participant’s perceptual threshold estimated after the initial training and inside the scanner. To determine this threshold, we employed a QUEST algorithm (Watson and Pelli, 1983) to estimate the Gabor tilt orientation for which participants’ performance was at 75%. Thresholds were determined when Gabors were superimposed with a grey circle. The scanning session started with the pre-conditioning phase (320 trials) employing an orientation discrimination task shown in **Figure 1A**. Participants were asked to indicate whether a Gabor stimulus was tilted clockwise or counter-clockwise relative to the horizontal meridian by pressing one of the 2 vertical buttons on a 4-button response pad (Current designs Inc., Philadelphia, PA) and to ignore the simultaneously presented task-irrelevant visual or auditory cues. The first scanning session terminated after the completion of pre-conditioning and participants attended the second session after at least 24 hours.

**Figure 1.**
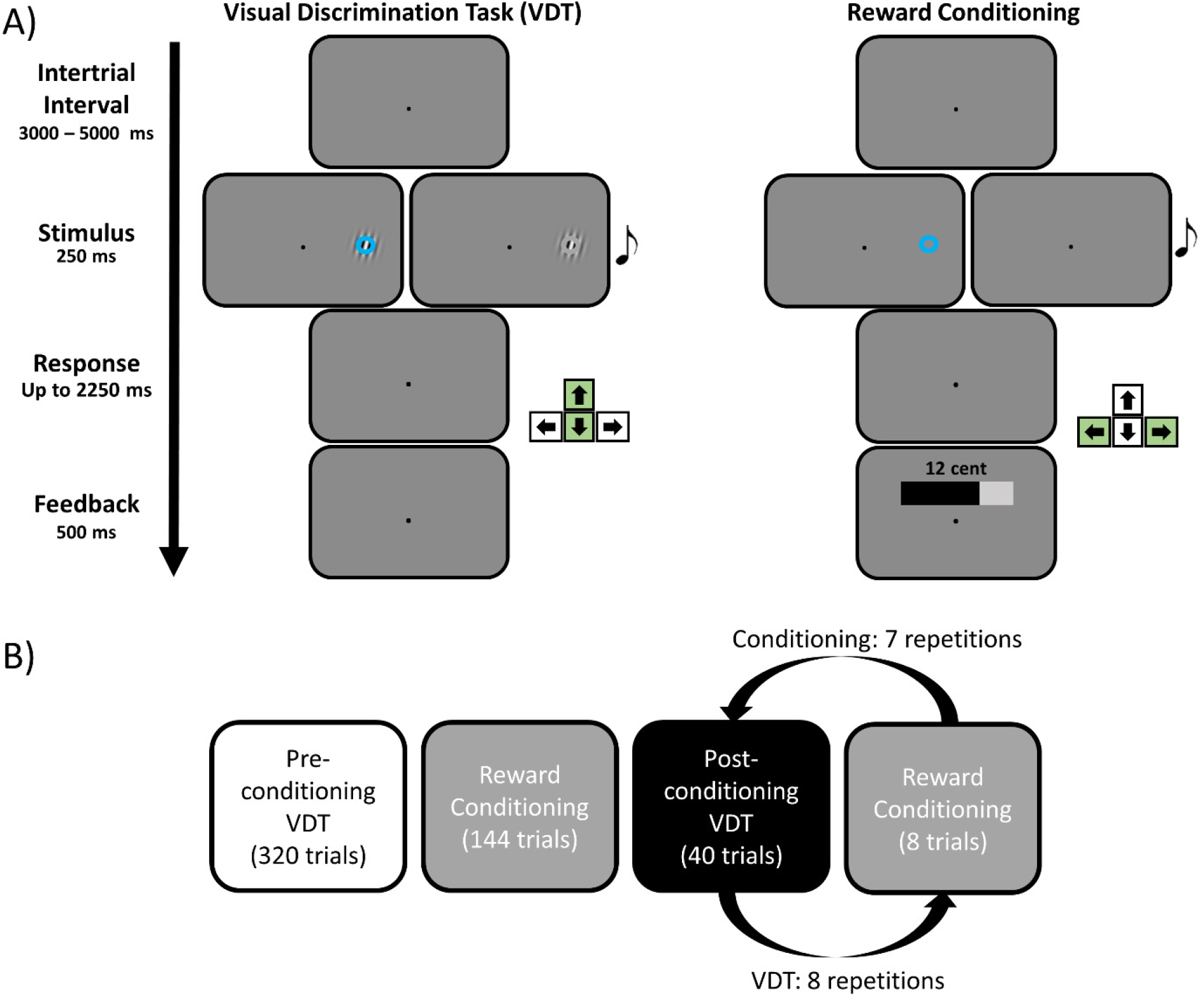
Behavioral paradigms. **A)** On the left side the visual discrimination task (VDT) used in the test phase is shown. Participants were instructed to discriminate the orientation of a Gabor patch (i.e. clockwise or counter-clockwise) overlaid with a semi-transparent ring by pressing upper or lower arrow keys on a response box, repectively. Simultaneously with the Gabor target, a task-irrelevant visual (intra-modal) or auditory (cross-modal) cue was also presented on the same location. The VDT task was employed both before and after a conditioing task (shown on the right side) where the reward associations of intra- and cross-modal cues were learned. During conditioing, participants were asked to indicate whether the cues (auditory or visual) were presented to the left or right side (by pressing the left or right arrow keys on a response box, repectively). The properties of the cues (color for the visual and pitch for the auditory tone) predicted different magnitudes of reward that was shown on the display during a feedback phase. During VDT, intra- and cross-modal cues were never predictive of reward delivery and accordingly the feedback display only contained a fixation point. **B)** The sequence of tasks employed during the experiment for each participant: first the VDT was completed before the cues were associated with rewards (referred to as the *pre*-conditioing phase recorded on day 1). Thereafter during the second session recorded on another day, participants first learned the reward associations of visual and auditory cues during the conditioing and then proceeded to the *post-*conditioning VDT with the cues that had been associated with rewards. To prevent the exinction of reward effects, the reward associations were reminded by interleaving the VDT with short conditioing blocks.

In the second scanning session, participants first completed a conditioning task to learn the reward associations of auditory and visual cues (see **Figure 1B**). During conditioning, participants were instructed to localize the visual (orange or blue rings) and auditory cues (pure tones 600 or 1000 Hz) and indicate whether they were presented to the left or to the right, by pressing one of the 2 horizontal buttons on a 4-buttons response pad. Upon correct response, participants saw the magnitude of the reward that was paired with a certain cue and thereby learned whether a visual or auditory stimulus was associated with high (mean = 25 Cents) or low (mean = 2 Cents, drawn from a Poisson distribution) monetary reward. All reward-cue combinations were counterbalanced across participants. In the third phase, referred to as post-conditioning (320 trials), the same procedure as in the pre-conditioning was employed with the exception that the task-irrelevant auditory and visual cues had already been associated with different amounts of monetary rewards. Additionally, in both pre- and post-conditioning one additional condition referred to as the neutral condition was included. The neutral condition contained the Gabor target overlaid by a semi-transparent grey ring. Since the grey color was never associated with any reward value during the conditioning, the neutral stimulus served as a means to measure target-specific responses in the visual cortex. Participants were instructed that they would get a bonus for each correct response, independent of the identity of the visual or auditory cues, though they would not be able to see the reward feedback.

In order to prevent extinction, we interleaved the post-conditioning blocks (each block with 40 trials) with a short conditioning block (8 trials). To ensure that participants had learned the reward-cue associations, we asked a question during and after the experiment. Based on these, all participants could correctly identify which cue properties were associated with high compared to low reward magnitudes.

### Behavioural data analysis

The data obtained from all parts of the experiment was analysed using custom-written scripts in MATLAB (version R2015a). For the behavioural analysis, we removed the trials in which participants did not respond or had extreme response times. To determine the extreme response times, we first log transformed each participant’s reaction times to achieve a roughly normal distribution and then removed trials which had reaction times >2SD from the mean (across all trials of each phase). This procedure removed 4.55, 4.67 and 5.13% of trials as outliers from the pre- and post-conditioning and conditioning, respectively. From the remaining trials, we calculated the mean of each response variable (accuracy and reaction times of correct trials) for each condition (high and low reward in auditory and visual cues) per subject during the post- and pre-conditioning separately. Afterwards, we entered the difference of accuracies and reaction times between the two phases (i.e., pre- and post-conditioning) as dependent variables in a 2×2 repeated measures ANOVA, with sensory modalities (intra-or cross-modal) and reward magnitude (high or low) as independent factors.

### MRI data acquisition

The imaging data was collected using Siemens Magnetom Prisma Fit (3T) with a 64 channels head coil at the University Medical Centre Göttingen. Structural images were acquired for each session using a MPRAGE T1-weighted sequence (FOV: 256 × 256mm; voxel size: 1 × 1 × 1mm; TR: 2250ms; TE: 3.3ms; number of slices: 176). Functional images were acquired using an EPI sequence (TR: 900ms; TE: 30ms; FOV: 210 × 210mm; voxel size: 3 × 3 × 3mm; slice thickness: 3mm; flip angle: 60°; number of slices: 45).

### fMRI data preprocessing

The imaging data was processed using the Statistical Parametric Mapping software (version SPM12: v7487; https://www.fil.ion.ucl.ac.uk/spm/). The data preprocessing pipeline consisted of realignment of the slices to the mean image, unwarping the images according to the voxel displacement mapped image, slice time correction for multiband interleaved sequence, coregistration between the functional and the structural images, segmentation of brain tissues according to the tissue probability maps, spatial normalization to the MNI space, and spatial smoothing with a kernel size of 8 mm (FWHM: 8 mm). All preprocessing steps were undertaken for the images that entered to the univariate GLM. For the multivariate analysis (MVPA), all steps were done except for the spatial normalization and spatial smoothing (see also under the *MVPA analysis*). For one participant the image required for unwarping could not be acquired due to technical problems at the scanner and we excluded this participant from all further fMRI-related analyses, resulting in N = 32 for corresponding results.

### Univariate GLM for effective connectivity

For the effective connectivity analysis, we designed a General Linear Model (GLM) with 10 regressors of interest for pre- and post-conditioning phases and 8 regressors of interest for the conditioning phase. These regressors were stick functions time-locked to the onset of the stimulus presentation in each trial (**Figure 1A**) and corresponded to the experimental conditions that varied in the reward magnitude (H-high or L-low), the sensory modality of the cues (V-visual or A-auditory), and the sides (L-left or R-right), plus the neutral trials (N: with no reward association) that was included in the pre- and post-conditioning in order to characterize the visual responses in the absence of reward associations.

Both scanning sessions were modelled in a single GLM, separated by a regressor marking session 1 (i.e. pre-conditioning phase) and session 2 (i.e. conditioning and post-conditioning phase). Moreover, we also modelled additional nuisance regressors such as six movement parameters for each session, three events of no interest (e.g. instructions) across the whole sessions, one regressor that marked the interleaved blocks of reward conditioning during the post-conditioing phase, and four regressors for marking each period of data acquisition (i.e. one regressor marked the pre-conditioning phase on day 1 and three regressors marked the three periods of data acquisition on day 2, one for conditioning and two for post-conditioning, each period corresponded to the time between the start and end of the scan).

### Multivariate analysis (MVPA)

For the MVPA analysis, we created a GLM where each trial in the pre- and post-conditioning was modelled as a separate regressor with stick functions at the onset time of the target. Four extra nuisance regressors were also included to model the inter-leaved blocks of conditioning and the instruction display, plus six head motion nuisance regressors. The trials during the conditioning blocks were modelled similarly as explained under *univariate GLM for the effective connectivity*. For this GLM, we used the images that underwent all preprocessing steps except for the spatial normalization and smoothing. The parameter estimates of this GLM (t values) were then fed into several pattern classifiers using LibSVM’s implementation of linear support vector machines (SVMs) (www.csie.ntu.edu.tw/~cjlin/libsvm). SVM classification was done using a whole-brain searchlight method, where the classification accuracy of each pattern classifier was computed based on the information contained in all voxels within a spherical searchlight region (radius: 6 mm) using a 10-fold cross-validation method. The searchlight was iteratively moved over every voxel in the whole-brain images and the calculated classification accuracy within each sphere was mapped to the voxel at the centre and normalized against the chance level accuracy (∼ 50% for a two-class pattern recognition). The output of the classifiers was used to compute first-level contrast images (see the description of orientation and value decoders below), which were then spatially normalized to the MNI space and smoothed (FWHM, 3 mm). These contrast images were then entered into a second-level analysis, in which the statistical significance of each contrast was evaluated using one-sample t tests.

Our pattern classification analysis comprised two main types of decoders: *an orientation decoder* to classify the tilt orientation of the target stimulus (i.e. classifying clockwise or counter-clockwise tilt orientation) and several *value decoders* to classify the associated reward magnitude of visual or auditory stimuli (i.e. classifying high or low reward magnitudes). These classifiers were designed to identify the early visual areas that encoded information about the target (orientation decoder) and brain regions that contained information about the associated reward value of stimuli (value decoders), respectively. Orientation decoders classified the stimulus orientation separately for different reward (high or low), cue modality (auditory or visual) and side (left or right). To examine the effect of reward value on early visual areas, we inspected the classification accuracy of this decoder using the contrast *High Value > Low Value* across all conditions (side and modality) during the post-conditioning corrected for the effects that existed prior to the learning of reward associations during the pre-conditioning. To identify the regions that contained information about reward value after learning of reward values, we built 2 types of value decoders: 1) value decoders that classified stimulus value across all conditions (i.e., both modalities: auditory or visual and locations\sides: left and right 2) value decoders that classified stimulus value separately for each sensory modality and each side. These decoders thus identified brain regions that were invariant to sensory modality and spatial location (*value decoder*_*1*_) or were sensitive to sensory modality and spatial location (*value decoder*_*2*_). The results of value decoders in post-conditioning were corrected against the results prior to the learning of reward associations in the pre-conditioning.

### Effective connectivity analysis (ECA)

In order to understand how cross- and intra-modal reward information is communicated across different brain regions to modulate early visual areas, we set up an effective connectivity analysis (ECA) using a dynamic causal modelling (DCM) approach (Friston et al., 2003; Friston, 2011). We hypothesized that there are three possibilities of how learned reward associations are communicated to modulate visual target processing. The first mechanism is based on a direct communication between the reward-related and the early visual areas, whereas the second mechanism relies on the involvement of either attention-related or sensory association areas to first process the reward information before it is further relayed to the early visual areas. Alternatively, reward-related information might be locally encoded in the early visual areas without the necessity of long-range communications across brain regions.

In order to test these hypotheses, we extracted the time series of regions of interest (ROIs) that were identified by our two types of MVPA decoders (i.e., orientation and value decoders) treating them as nodes in DCM networks to be modelled. Both types of decoders could potentially identify multiple brain regions (see the Results and **Table 2**). Therefore, we limited our analysis to ROIs that were most informative for testing our a priori hypotheses. These ROIs comprised the early visual areas (EVA) known to contain information about the stimulus orientation (Hubel and Wiesel, 1968; Grill-Spector and Malach, 2004) and valuation areas that based on previous literature are known to play a role in coding stimulus value and attentional or sensory processing. The visual ROIs (see **Table 2, Figure 2B** and **Supplementary Figure 2**) were defined as regions that had a significantly higher orientation classification accuracy in the presence of high compared to low reward stimuli across both modalities (i.e. the contrast: *High Value > Low Value*) in post-compared to pre-conditioning and were within an anatomical mask consisting of bilateral V1-V2 areas (Eickhoff et al., 2005). In order to define the ROIs that contained information about the stimulus associated value, we inspected the results of our two *value decoders* (see also the description of MVPA methods). The classification results of *value decoder*_*1*_ revealed a right lateralized inferior orbitofrontal area ([51 26 −7], *p* uncorrected < .005, k = 20), an area known to encode stimulus associative value (Kringelbach, 2005; Zald et al., 2014). The output of the *value decoder*_*2*_ was inspected either across sensory modalities or based on an interaction contrast that tested whether a region contained more information about the associated value of a specific sensory modality over the other (e.g. classification accuracy is higher for auditory than visual). Among the activations revealed by the first contrast (**see Table 2**), we selected both the strongest cluster at the right superior temporal areas (at [57 −28 8], *p* uncorrected < .005, k = 20), an area related to multisensory processing (Calvert et al., 2000; Stein and Stanford, 2008) and has been observed to be modulated by reward magnitude (Pooresmaeili et al., 2014), and the largest cluster that corresponded to the left anterior intraparietal sulcus (IPS) (at [−33 −58 53], *p* uncorrected < .005, k = 20), an area known to play a role in the allocation of attention (Corbetta et al., 2000; Corbetta and Shulman, 2002; Serences and Yantis, 2007) and has well-documented neuroanatomical connections with the frontal areas (Greenberg et al., 2012).

**Figure 2.**
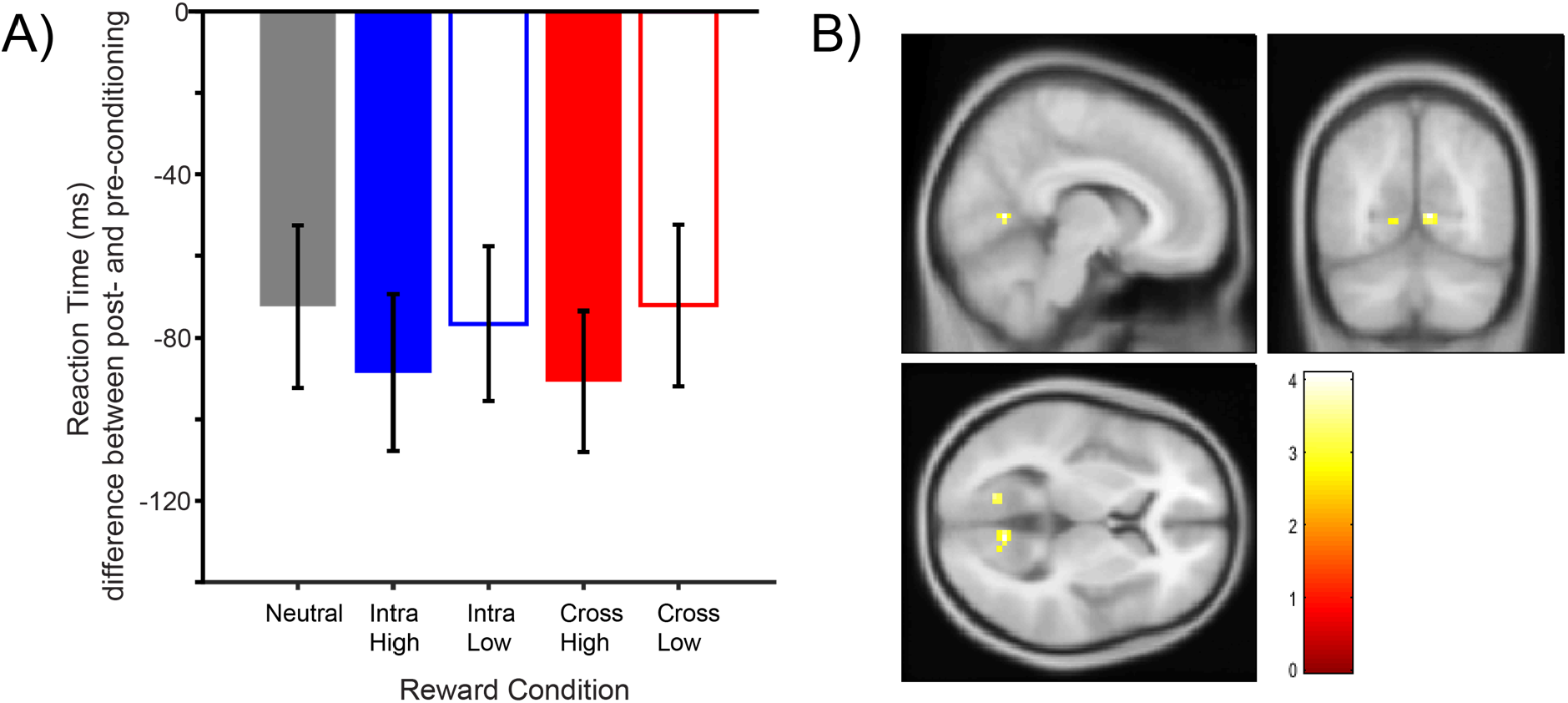
Behavioral and BOLD effects of reward on visual discrimination. **A)** Baseline corrected reaction times for all conditions. Error bars are s.e.m. **B)** Reward facilitation in early visual areas (masked with V1-V2 anatomical mask from Eickhoff and colleagues (2005)). The activations correspond to regions in area V1-V2 where the classification accuracy of the orientation decoder was higher for high compared to low reward condition during the post-conditioning after correcting for differences in pre-conditioning. Activations are shown at an uncorrected p < .005, k = 10, revealing a peak in the right hemisphere located at xyz *=* [9 −64 5] and in the left hemisphere at xyz = [−12 −67 2].

For each ROI, time series were extracted separately for pre- and post-conditioning by overlaying the group functional ROI on each participant’s structural scan. Within this framework, we estimated 11 biologically plausible models for the pre- and post-conditioning phases in which the directed causal influences among brain regions could change by three types of parameters: driving inputs and intrinsic and modulatory connections. Driving inputs corresponded to the incoming visual information contained in the different experimental conditions. To estimate the driving inputs, we used the univariate GLM which provided us the estimated BOLD times series corresponding to our 5 experimental conditions: intra-modal high reward (VH), intra-modal low reward (VL), cross-modal high reward (AH), cross-modal low reward (AL), and neutral (N). For each input the data of two sides (left and right) were combined and fed to the DCM models. Furthermore, as all stimuli contained the same visual target (i.e. the Gabor patch), we fed all driving inputs to the visual ROI (EVA) which is the first stage of information processing in a visual task. Intrinsic (condition-independent) connections were defined between every pair of nodes in the network and as self-connections. The models differed from each other with respect to the modulatory connections, which varied with the experimental conditions (**Figure 4**). In the null model, only intrinsic connections were included and no condition-dependent modulatory connection existed. The rest of the models assumed different patterns of connectivity between the early visual areas and other ROIs. One class of models (model 1-4) assumed that the valuation ROI (i.e. lateral OFC) communicated with the early visual areas indifferently across intra- and cross-modal condition. Specifically, the communication might involve a long-range direct communication (model 1), where previous studies have shown that lateral OFC receives direct inputs from visual and auditory cortices (Kringelbach and Radcliffe, 2005), making the assumption of a direct connection between the two regions plausible. Another possibility was that the communication of the valuation and visual ROIs is indirect, with the information being first relayed to sensory-related ROI for cross-modal condition (model 2). Specifically, these models involved a modulatory connectivity between OFC-STS (Zald et al., 2014) and thereafter from STS to EVA (V1-V2 areas), comprising connectivity patterns that are supported by previous studies (Felleman and Van Essen, 1991; Lewis and Noppeney, 2010). The third possibility was that the valuation and visual ROIs influenced each other through engaging the attention-related areas, i.e. IPS in our case; (model 3) or both attentional and sensory areas (model 4). The pattern of inter-areal connectivity assumed by these models is in line with previous literature showing functional and structural connectivity between these areas: lateral OFC is functionally connected with IPS (Zald et al., 2014), IPS has connections to STS as demonstrated by diffusion tractography (Bray et al., 2013) and IPS has structural connections to early visual areas (Felleman and Van Essen, 1991; Bray et al., 2013). Moreover, STS has been known to have a functional connection with the primary visual area (Noesselt et al., 2007). So far, model 1-4 assumed that intra- and cross-modal cues behaved similarly. In order to capture the possibility of a dissociation between intra- and cross-modal pathways, we also modelled another class of models (model 5-10) where distinct pathways were involved in intra- and cross-modal reward processing. Lastly, we also included a *null* model (model 11), which assumed that the influence of reward on early visual areas occurred locally with these areas and did not require a constant log-range communication with other areas.

These models were therefore captured by a DCM model space consisting of 11 models per phase (pre-or post-conditioning). Each model was estimated for each participant and each phase (pre- and post-conditioning) separately. Then, models were compared using a group-level random effects Bayesian Model Selection (BMS) approach (Stephan et al., 2009) to select the most probable model given the observed BOLD time-series. We employed a random effect (RFX BMS) to select the winning model, as this method allows for the possibility that different participants may have different preferred models. Note that in all models (see **Figure 4**), high and low reward conditions in both phases are assumed to be processed by the same brain regions and involve the same inter-areal connectivity patterns, albeit the strength of these connections were hypothesized to differ depending on the reward magnitude (between phases: pre- and post-conditioning and reward conditions: high and low). To test this latter hypothesis, we next inspected the winning model detected by BMS approach and tested whether the connectivity strength of this model was modulated by reward magnitude using a Parametric Empirical Bayes (PEB) approach (Zeidman et al., 2019). The PEB approach is a hierarchical Bayesian model that uses both non-linear (first-level) and linear (second-level) analyses. The advantage of using this approach is that inter-individual variability in model parameters is parameterized at the second level. Hence, parameter estimates for subjects with noisy data are likely to be adjusted in order to conform to the group distribution. Combining BMS and PEB approaches allowed us to maximally capture the inter-individual variability while selecting models using BMS, while having a more sensitive measurement of parameter estimates of the winning models by using PEB that adjusts parameter estimates based on their distribution across the participants. As our model comparison analysis revealed that model 10 had the strongest evidence in the post-conditioning, while the *null* model had the strongest evidence in the pre-conditioning, we exclusively extracted the parameters of the winning model 10 for pre- and post-conditioning of each participant as the input of the design matrix in the PEB. Then, at the group level, we constructed a PEB model with a constant term (mean parameter estimates across participants) and an additional binary regressor to model the difference between pre- and post-conditioning. This allowed us to investigate how the connectivity strength was modulated by reward magnitude before and after participants had learned the reward-cue associations. As we were interested in the reward modulation of each connection between regions, we focused on the estimated parameters in the modulatory (i.e. B matrix) connectivity, specifically for both feedforward/bottom-up and feedback/top-down connections. Finally, for each connection, we report the reward modulation (high-low) posterior probabilities using a threshold of > 0.99, correcting for multiple comparisons across connections (Bonferroni correction).

## Results

### Conditioning phase: Recruitment of the classical brain regions involved in the reward associative learning

Participants exhibited near perfect accuracy in localizing both visual and auditory stimuli (both > 95%), however there was no significant effect of reward on either the response accuracies or the reaction times (for details see the Supplementary Information). Analysis of the BOLD responses revealed the classical brain areas that are involved in the associative learning of rewards, such as the ventral striatum and insula (see the **Supplementary Figure 1**). The effect of reward on the BOLD responses was largely independent of the sensory modality, except for the higher activations observed for the auditory compared to visual reward value found in the right caudate (see the **Supplementary Table 1**).

### Previously reward-associated cues slightly enhanced the speed of visual discrimination during the post-conditioning

We next examined the behavioural effects of rewards from the same (intra-modal) or different (cross-modal) sensory modality on the visual discrimination task. Compared to the pre-conditioning, reaction times decreased for all conditions during the post-conditioning phase indicating that with longer training on the task, participants’ speed of perceptual decisions increased (**Table 1** and **Figure 2A**). This speed enhancement was stronger for the high compared to low reward conditions. Accordingly, we found a main effect of reward on the reaction times as higher reward magnitude increased the speed of visual discrimination across sensory modalities (F(1,32) = 4.46, *p* = 0.04, η_p_^2^ = 0.12). Other main and interaction effects did not reach statistical significance. The effect of reward in individual conditions (cross- and intra-modal conditions) was not significant (both ps>0.1), and although high reward stimuli seemed to lead to faster responses compared to the neutral condition, this effect did not reach statistical significance (F(2,64) = 1.34, *p* = 0.268, η_p_^2^ = 0.040). Analysis of the accuracies revealed neither a main effect of reward value nor an interaction with the sensory modality (both Fs<1.5 and ps>0.1). Together, these results indicate a weak behavioural advantage for high compared to low reward stimuli in our experiment which was mainly observed for the reaction times.

**Table 1.** Behavioral results during the visual discrimination task performed in pre- and post-conditioning

| Condition | RT<br>(pre-conditioning) | Accuracy<br>(pre-conditioning) | RT<br>(post-conditioning) | Accuracy<br>(post-conditioning) |
| --- | --- | --- | --- | --- |
| High Reward Intramodal<br>(HV) | 938.33±24.13 ms | 81.08±1.09% | 849.72±20.25 ms | 80.53±1.33% |
| Low Reward Intramodal<br>(LV) | 929.56±23.77 ms | 80.80±1.29 % | 852.98±19.71 ms | 80.23±1.48% |
| High Reward Cross-modal (HA) | 934.15±26.25 ms | 82.25±1.07% | 843.40±20.79 ms | 82.25±1.38% |
| Low Reward Cross-modal (LA) | 920.64±26.08 ms | 84.48±1.71 % | 848.50 ±21.59 ms | 84.48±1.36% |
| Neutral | 925.08±24.04 ms | 80.68±1.40% | 852.67±19.71 ms | 80.66±1.60% |

### Reward-driven modulation of target representations in the early visual areas during the post-conditioning

We next examined how the reward value affected the encoding of the target’s tilt orientation in the early visual areas. To this end, we examined the results of the whole-brain searchlight *orientation decoder* (for classification of clockwise and counterclockwise orientations) and identified areas within an anatomical mask of area V1-V2 which exhibited a reward-driven increase in the decoding accuracy across sensory modalities in the post-compared to the pre-conditioning.

This contrast revealed a bilateral activation with a peak at xyz = [9 −64 5] on the right and at xyz = [−12 −67 2] on the left visual cortex (**Figure 2B**). Importantly, this activation overlapped with the regions within areas V1-V2 that were activated by the Gabor stimulus in the neutral condition indicating that they corresponded to the target-specific representations within the early visual areas (**Supplementary Figure 2**). This result indicates that higher reward enhanced the neural representation of the visual target already as early as in area V1-V2, in line with previous findings where reward-driven enhancement of the magnitude (Serences, 2008) or the specificity of spatial patterns (Pooresmaeili et al., 2014) of neural responses were observed in the early visual areas. Additionally, to further support this finding, we checked the opposite contrast (classification accuracy in Low Value > High Value) using the same threshold and mask, and did not find any activation.

After establishing that higher reward enhances the reliability of target representations in the early visual areas, we asked *where* in the brain the associated reward value of stimuli is encoded and *how* the reward-related signals are communicated to visual areas. In order to answer these questions, we conducted two types of analyses: 1) An MVPA analysis to identify *where* in the brain the reward value is encoded, and 2) An effective connectivity analysis in which the possible communication patterns between the identified valuation regions and early visual areas were tested (thus answering the question of *how*).

### Identification of the brain regions that encode stimulus value during the post-conditioning (*where*)

Towards answering the first question regarding *where* in the brain the stimulus value is encoded after learning of the reward associations, we inspected the results of our two value decoders. To identify brain areas that are responsive exclusively to the stimulus reward magnitude irrespective of its sensory features (sensory modality and location), we inspected the results of the *value decoder 1* (see Material and Methods). This decoder performed a whole-brain search for regions that contained information about the reward value after value associations were learned (class labels were: high or low reward magnitude, see Material and Methods). The classification accuracy of *value decoder 1* was highest in a cluster in the left orbitofrontal cortex (blue cluster in **Figure 3, Table 2**, and **Supplementary Figure 3**), while several other areas related to the reward processing such as ventral striatum, ventromedial prefrontal cortex were also identified by this analysis (**Table 2**). The lateral OFC cluster was further selected for the subsequent effective connectivity analysis.

**Table 2.** Whole-brain activations of value decoders thresholded at uncorrected p < .005 and k = 20. Regions marked with bold font were selected as ROIs used for the effective connectivity analysis.

| Cluster size | MNI coordinates (in mm) |  |  | T | p | Side | Region |
| --- | --- | --- | --- | --- | --- | --- | --- |
|  | x | y | z |  |  |  |  |
| Results of *Value Decoder 1*: areas that distinguish between high and low value irrespective of sensory properties |  |  |  |  |  |  |  |
| 43 | 51 | 26 | -7 | 6.08 | 0.006 | R | Inferior orbitofrontal |
| 44 | -45 | -46 | -49 | 5.27 | 0.006 | L | Cerebelum |
| 80 | 36 | -79 | -52 | 4.69 | 0.000 | R | Cerebelum |
| 37 | 42 | -61 | -4 | 4.39 | 0.01 | R | Inferior temporal |
| 22 | 3 | 4 | 11 | 4.06 | 0.038 | L | Caudate |
| 23 | 12 | 8 | 41 | 3.64 | 0.034 | R | Cingulate cortex |
| 22 | -42 | 23 | -16 | 3.55 | 0.038 | L | Inferior orbitofrontal |
| 23 | -3 | 65 | -7 | 3.52 | 0.034 | L | Medial orbitofrontal |
| Results of *Value Decoder 2*: areas that distinguish between high and low value for each location and sensory modality. After value classification was performed, results were inspected across sensory modalities. |  |  |  |  |  |  |  |
| 37 | 57 | -28 | 8 | 4.62 | 0.01 | R | Superior temporal |
| 34 | -6 | -73 | 23 | 4.37 | 0.012 | L | Cuneus |
| 36 | 9 | -37 | 44 | 4.35 | 0.01 | R | Cingulate cortex |
| 69 | -33 | -58 | 53 | 4.03 | 0.001 | L | Inferior parietal |
| 28 | -18 | -52 | 8 | 4.00 | 0.021 | L | Precuneus |
| 20 | -51 | 44 | -1 | 3.82 | 0.046 | L | Inferior orbitofrontal |
| 39 | -12 | -25 | 71 | 3.80 | 0.008 | L | Motor cortex |
| 24 | 21 | -28 | 53 | 3.54 | 0.031 | R | Somatosensory |
| 22 | -54 | -55 | 26 | 3.40 | 0.037 | L | Temporoparietal |
| 23 | -57 | 2 | -1 | 3.28 | 0.034 | L | Temporal pole |
| Results of *Value Decoder 2*: areas that distinguish between high and low value for each location and sensory modality. After value classification was performed, results were inspected for intra-modal stimuli. |  |  |  |  |  |  |  |
| 27 | -12 | -22 | 71 | 4.81 | 0.024 | L | Paracentral lobule |
| 20 | 57 | -31 | 8 | 4.72 | 0.047 | R | Superior temporal |
| 23 | 45 | -28 | 53 | 4.58 | 0.035 | R | Postcentral |
| 36 | -36 | -61 | 56 | 4.49 | 0.011 | L | Superior parietal |
| 32 | -24 | 23 | 56 | 4.45 | 0.015 | L | Frontal mid |
| 28 | 3 | 50 | 32 | 4.12 | 0.022 | R | Frontal sup medial |
| 26 | 9 | -55 | -43 | 3.91 | 0.026 | R | Cerebellum |
| 20 | -36 | -76 | -1 | 3.68 | 0.047 | L | Occipital mid |
| 21 | -9 | -82 | -28 | 3.54 | 0.043 | L | Cerebellum |
| 22 | -51 | -64 | -16 | 3.46 | 0.039 | L | Inferior temporal |
| **Results of Value Decoder 2:** areas that distinguish between high and low value for each location and sensory modality. After value classification was performed, results were inspected for **cross-modal stimuli**. |  |  |  |  |  |  |  |
| 60 | -39 | -25 | 5 | 5.17 | 0.001 | L | Heschl |
| 37 | 54 | -28 | 5 | 4.66 | 0.008 | R | Superior temporal |
| 20 | 48 | -1 | 2 | 4.56 | 0.041 | R | Insula |
| 30 | -6 | -73 | 23 | 4.49 | 0.015 | L | Cuneus |
| 25 | -66 | -28 | 17 | 4.35 | 0.024 | L | Superior temporal |
| 41 | -57 | -55 | 29 | 4.24 | 0.006 | L | Angular |
| 66 | 12 | -25 | 38 | 4.07 | 0.001 | R | Cingulum mid |
| 24 | -42 | 5 | -19 | 3.77 | 0.027 | L | Temporal pole sup |
| 27 | -15 | -46 | -13 | 3.69 | 0.02 | L | Fusiform |
| 21 | -18 | -52 | 8 | 3.44 | 0.037 | L | Calcarine |
| **Results of Value Decoder 2:** areas that distinguish between high and low value for each location and sensory modality. After value classification was performed, results were inspected for the interaction of **cross-modal>intra-modal**. |  |  |  |  |  |  |  |
| 36 | -42 | -19 | 5 | 3.78 | 0.009 | L | Heschl gyrus |
| **Results of Value Decoder 2:** areas that distinguish between high and low value for each location and sensory modality. After value classification was performed, results were inspected for the interaction of **intra-modal>cross-modal**. |  |  |  |  |  |  |  |
| **No voxel survived** |  |  |  |  |  |  |  |

**Figure 3.**
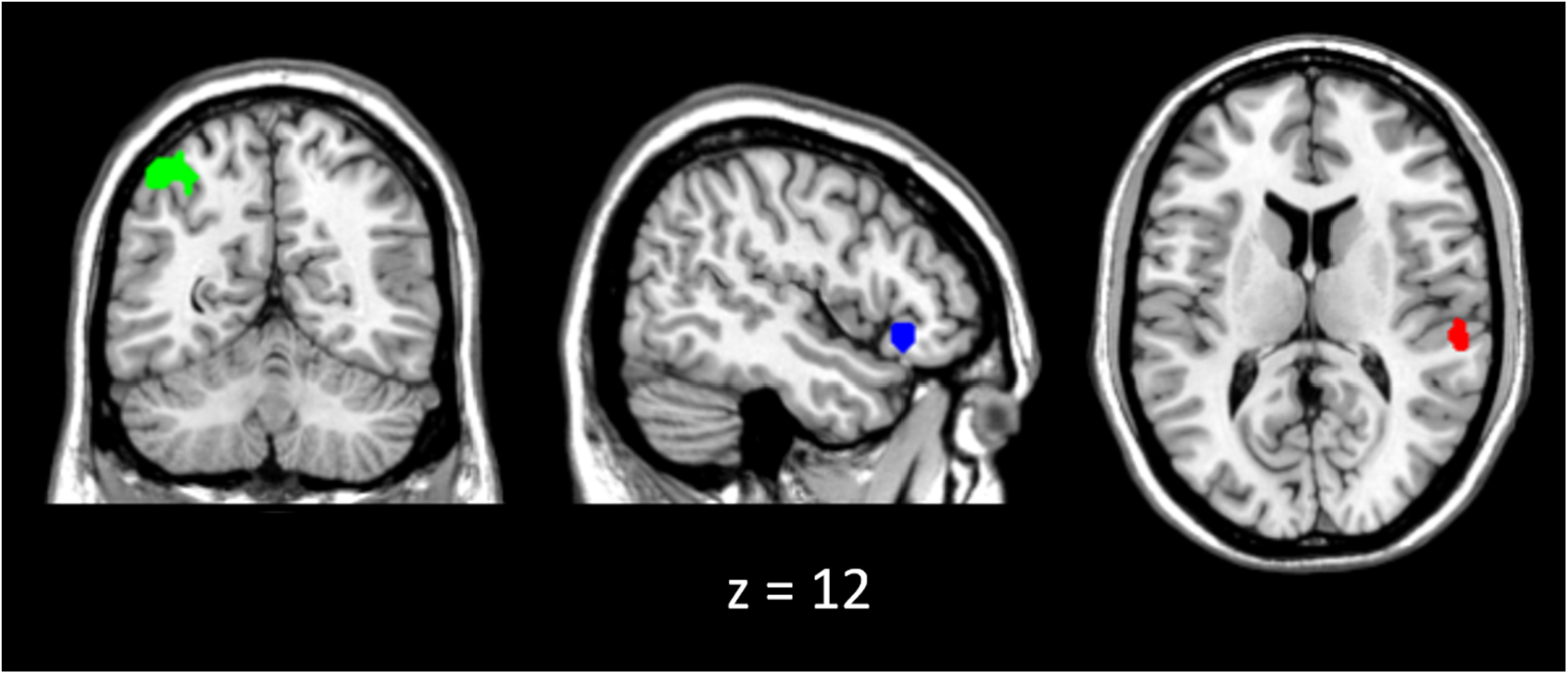
Regions of interest identified by the *value decoders* and used for the effective connectivity analysis. *Value decoder1* identified a cluster in the OFC xyz = [51 26 −7] shown in blue, which discriminated high and low value stimuli irrespective of their sensory properties (i.e., location and sensory modality). *Value decoder2*, classified high and low reward stimuli from each location and sensory modality separately and showed clusters in IPS xyz = [−33 −58 53] in green and STS xyz = [57 −28 8] in red, where reward value was reliably decoded across sensory modalities. The activations are shown at uncorrected *p* < .005 with k = 20, and the cursor is located at xyz = [48 −58 12] to illustrate all ROIs.

Next, we asked which brain areas are involved in the encoding of stimulus value specifically for each sensory modality and stimulus location. These areas are instrumental in conveying additional information regarding the specific sensory feature of reward cues across the brain. In order to investigate this question, we examined the results of the *value decoder 2* which decoded the stimulus value separately for each sensory modality (intra- and cross-modal) and stimulus location (left and right, see the Material and Methods). We then inspected the results of this decoder across both sensory modalities as well as differentially contrasting one modality against the other. The strongest reward modulation across sensory modalities was observed in the superior temporal areas (STS, red cluster in **Figure 3** and **Supplementary Figure 3**), an area that is tightly linked with the multisensory processing (Calvert et al., 2001; Stein and Stanford, 2008). Interestingly, we also found that across sensory modalities stimulus value was reliably decoded from regions with a known role in attentional processing such as a large cluster in the anterior intraparietal area (IPS, **Figure 3** and **Supplementary Figure 3**). This area has not only been related to the attentional selection (Corbetta and Shulman, 2002), but also has been shown to be modulated by reward (Platt and Glimcher, 1999; Bendiksby and Platt, 2006; Louie et al., 2011). Moreover, we also observed several areas such as the cuneus, cingulate, temporoparietal area, and also motor cortex which contained reliable representations of stimulus value across modalities (see **Table 2**).

To test whether there are specific brain areas that contain more information about the stimulus value from one compared to another sensory modality, we contrasted the whole-brain results of the *value decoder 2* for Auditory (Cross-modal) >Visual (intra-modal) and vice versa. The first contrast (i.e. classification accuracy in auditory > classification accuracy in visual), revealed a cluster in the left auditory cortex which corresponded to the primary auditory area (area A1, at *p* < 0.005, k = 20 uncorrected). However, in the intra-modal interaction (i.e. classification accuracy in visual > classification accuracy in auditory), no voxel survived at the same threshold (at *p* < 0.005, k =20 uncorrected, see **Table 2**).

Based on the above results and our a priori hypotheses, we took the IPS and STS clusters as ROIs that might be involved in the long-range communications between the valuation network (i.e., OFC, identified by value decoder 1) and the early visual areas (i.e, EVA, identified by the orientation decoder), as they were discriminative of reward value across sensory modalities. Furthermore, value decoder 2 only identified the primary auditory cortex (area A1) as an area that contained more information about one over the other sensory modality (cross-modal > intra-modal), whereas we did not find any area that selectively encoded the value of intra-modal stimuli. In contrast to the A1, that might play a role in processing the sensory features of the auditory reward-associated cues, the superior temporal areas are known to be involved in higher-order auditory processing and the integration of information across senses (Stein and Stanford, 2008), where most likely both the visual target and auditory reward-associated cues were processed. In fact, when we inspected the results of value decoder 2 in each individual modality, we observed STS activations for both intra- and cross-modal value (**Table 2**). We therefore reasoned that including STS but not A1 in our effective connectivity analyses would capture the reward-driven effects of both cross-modal and intra-modal stimuli, while reducing the complexity of models by adding multiple areas with overlapping functionalities (i.e., STS and A1).

### Effective connectivity analysis revealed *how* reward information is broadcasted across the brain

After identifying the potential brain areas that might mediate the reward-driven modulation of early visual areas, we tested possible models of *how* reward information is broadcasted across the brain using an effective connectivity approach. Based on our hypotheses, three possibilities existed which gave rise to 11 biologically plausible schemes in our model space (**Figure 4A**): **1)** reward signals are communicated indifferently from the reward-related areas to the early visual areas, involving either a long-range direct projection (**fig.4A**, model 1) or mediation through the attention-related or higher sensory-related areas (**fig.4A**, model 2-4), **2)** reward signals are communicated following a modality-specific pathway through attention and/or higher sensory-related areas (**fig.4A**, model 5-10), or **3)** reward signals have a long-lasting effect where the neural plasticity in the early visual areas is altered locally without the necessity of information flow from and to the other brain areas (**fig.4A**, model 11 or *null*, see the Material and Methods). These models thus differed with respect to the nodes/regions and connectivity patterns which underlay the intra-modal and cross-modal information transfer. In all models, high and low reward conditions involved the same nodes and connectivity patterns but could influence the strength of the connectivity between each pair of nodes to a different extent (see Material and Methods). Therefore, we first established which nodes and connectivity patterns best explained the BOLD times series of the intra- and cross-modal conditions in pre- and post-conditioning and thereafter tested whether the strength of connections in the winning model was modulated by reward magnitude after the stimulus-reward associations were learned.

**Figure 4.**
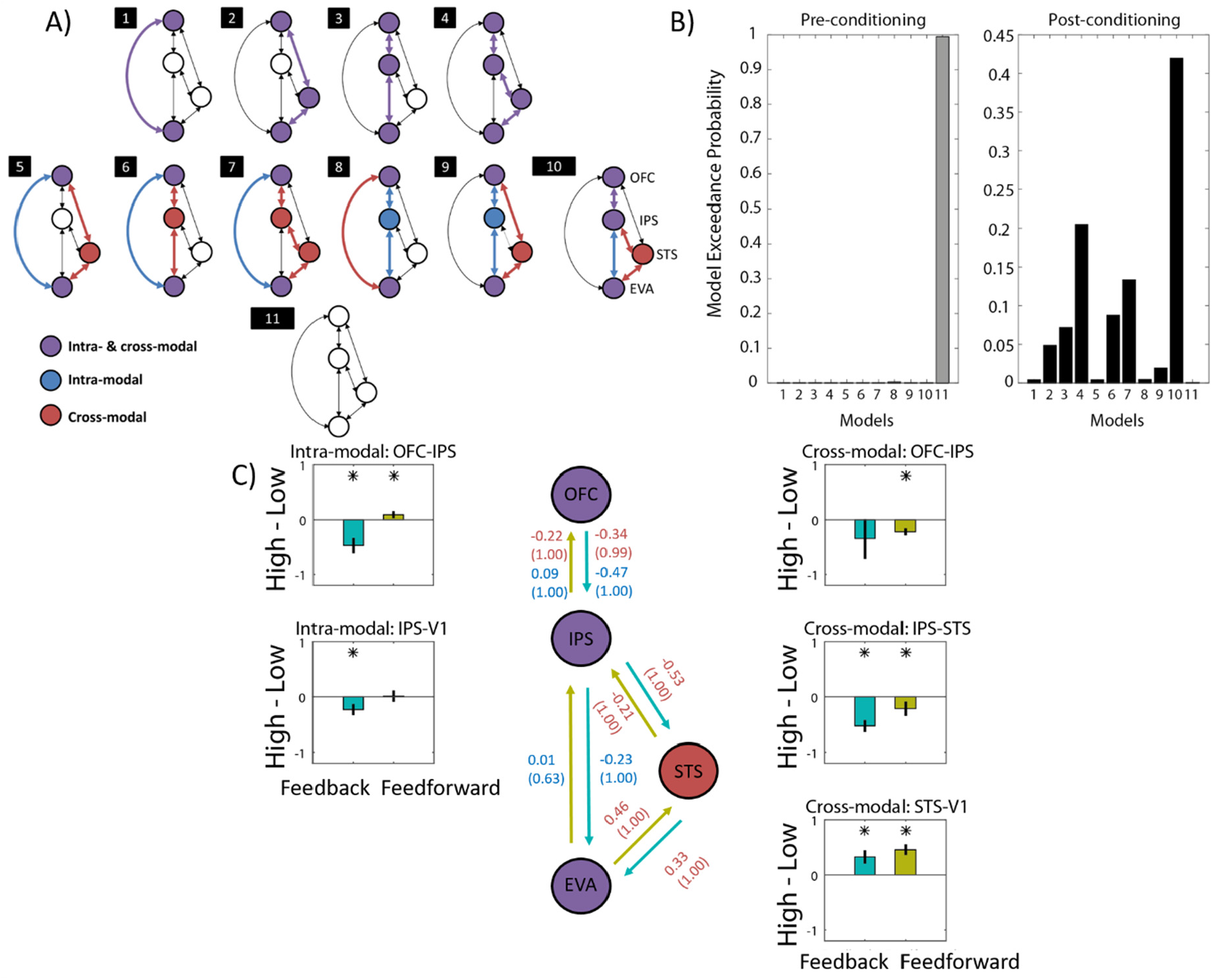
Effective connectivity results. **A)** Schematic of 11 models that were considered to probe the mode of the bidirectional communication between the reward-related areas and the early visual areas (EVA). **B)** The models were estimated for both pre-(in grey) and post-conditioning (in black) phases. The exceedance probabilities of random effects Bayesian model selection demonstrated that model 11 (null model) wins in pre- and model 10 wins in post-conditioning. **C)** Estimated parameters (in Hz) of the winning model in post-conditioning were used to characterize the reward modulation (i.e. changes in the strength of each connection when comparing high relative to low rewards) corrected for effects before reward associations were learned (i.e. post – pre conditioning). Reward modulations are shown for each connection between two regions and separately for each direction (feedback and feedforward, in teal and dark yellow, respectively). * corresponds to *p* < 0.01 (equivalent to posterior probabilities > 0.99) and corrected for multiple comparison using Bonferroni correction. Errorbars depict 99% confidence intervals of the subtracted distribution (high – low). The middle panel illustrates the schematic of the winning model and depicts the strength of reward modulation for feedforward and feedback connections (teal and dark yellow arrows, respectively) and their respective posterior probability (in bracket) for the intra-modal (blue) and cross-modal (red) conditions.

Among the possible models, our results (**Figure 4B**) indicated that model 10 gained the highest evidence in the post-conditioning (*p*_ex = 0.42) relative to the second best model (model 4, *p*_ex = 0.2). Meanwhile, model *null* gained the highest evidence in the pre-conditioning (*p*_ex = 0.99). As expected, learning of the reward associations changed the way that information was communicated across the brain, as reward-related areas were only involved in modulating the early visual areas *after* the stimulus-reward associations had been established. In the winning model 10 in the post-conditioning, intra- and cross-modal information needed to be gated through the regions involved in the attentional selection, as IPS was involved in mediating both communication paths. Additionally, the cross-modal condition engaged the STS, a higher-order sensory area, in order to communicate the reward information across the brain. This is aligned with our hypothesis 2, where intra- and cross-modal effects were mediated through both attention and sensory-dependent areas.

In order to infer how reward value modulated the strength of connectivity between every pair of nodes/regions in the winning model, we next conducted a group level analysis on the weights of feedforward and feedback connections. We included both pre- and post-conditioning data of the winning model (model 10 in the post-conditioning) in our design matrix and examined the reward-driven changes in the weights of connections that occurred after the stimulus-reward associations were learned by regressing out the effects in the pre-conditioning (see the Material and Methods). This analysis summarised in **Figure 4C**, revealed widespread effects of reward value on the strength of connections between different regions. Specifically, we found both supra-modal (modality-independent) and modality-dependent reward modulations. The feedback from the valuation area (OFC) to the mediation areas in the IPS in intra-modal and STS in cross-modal condition were regulated similarly (i.e. modality-independent), as in both cases a feedback inhibition was observed (OFC-IPS: −0.47 Hz and −0.34 in intra- and cross-modal, respectively; and IPS-STS in cross-modal: −0.53Hz), likely to prevent the allocation of processing resources to high reward cues that were irrelevant to the target discrimination. However, there was a dissociation in the feedforward communication paths (i.e. modality-dependent), where intra-modal cues relied on the excitatory modulation (IPS-OFC: 0.09 Hz) and cross-modal cues relied on the inhibitory modulation (STS-IPS: −0.21 Hz and IPS-OFC: −0.22 Hz). This dissociation between intra- and cross-modal feedforward connections might indicate that mediation areas (IPS and STS) engage distinct mechanisms to prioritize the processing of sensory features of the high reward stimuli. Specifically, feedforward processing of intra-modal rewards was enhanced due to the need to discriminate the intra-modal reward cues from the visual target as both emanated from the same sensory modality, whereas the feedforward processing of cross-modal reward cues that were distinct from the visual target decreased. Moreover, the dissociation of reward effects was further observed in the communication between the mediation areas and the early visual areas (EVA), where intra-modal cues relied more on the inhibitory and cross-modal cues on the excitatory feedback modulation. Specifically, whereas the feedback communication in the intra-modal condition was suppressed (−0.23 Hz), both feedback (0.33 Hz) and feedforward (0.46 Hz) communication paths were facilitated for cross-modal cues. This distinction might indicate that the way higher reward increases the perceptual discriminability of the target may differ between the intra- and cross-modal conditions, where intra-modal rewards boost the differentiation and cross-modal rewards increase the integration of the reward cues and the target. Accordingly, the top-down inhibitory modulation from the IPS to EVA likely suppressed the processing of the high reward intra-modal cues (i.e. irrelevant information) to improve the representation of the target. In contrast, enhancing the feedforward processing of the visual target in EVA-STS, could potentially enhance the integration of the auditory reward-associated cues and the visual target and subsequently the excitatory feedback from the STS to EVA could boost the representation of the target.

## Discussion

This study aimed to investigate the reward-driven modulation of the early visual processing. We compared intra- and cross-modal previously reward associated cues to probe whether their reward-driven effects depended on the sensory modality of the cues. In our paradigm using a visual discrimination task, previously reward associated task-irrelevant cues slightly improved the speed of perceptual decisions. Moreover, using a multivariate pattern classification approach, we observed that high reward stimuli enhanced the neural representations of the target in the early visual areas. We looked further into the possible neural mechanisms governing this effect by means of an effective connectivity analysis. This analysis revealed that reward-related information is communicated across the brain in both modality-independent and modality-dependent manners. In general, the reward-driven effects of both intra- and cross-modal cues recruited areas involved in the encoding of reward value and attentional selection. However, cross-modal rewards additionally involved the higher-order sensory-related areas such as STS. The feedback communication between these areas was predominantly inhibitory, demonstrating that reward value modulates the prioritization of information processing. Unlike the modality-independent interactions observed between the higher-level areas, the neural communication to and from the early visual areas were differentially modulated by intra- and cross-modal rewards. At this level, intra-modal rewards produced predominantly feedback inhibition whereas cross-modal rewards led to excitatory feedforward and feedback modulations.

Previously reward associated cues have been known to capture attention (Anderson et al., 2011). Consequently, when reward cues are not the target of the task, response times are slowed down as attention needs to be re-oriented from the reward-associated task-irrelevant distractors to the target. In our study, we observed a weak facilitation (i.e. faster reaction times) by the irrelevant high reward cues. A possible reason is the spatial alignment of the reward cues and target in our study that differed from Anderson and colleagues (2011), where in their design, reward cues and target were separated spatially. In contrast, in our design reward cues and the visual target were presented at the same location. Therefore, attention did not need to be re-oriented and the capture of attention created by the irrelevant reward cues could potentially spill over to the target, energizing the responses. Moreover, in contrast to our previous study (Vakhrushev et al., 2021), where perceptual discrimination and visual evoked potentials were either suppressed or enhanced by the intra- and cross-modal rewards, respectively, we did not observe an interaction effect. An aspect that differed with this previous study was the length of training on the task before the reward associations were learned, where in the current study the number of trials in the pre-conditioning phase was doubled so that participants are better accustomed to the reward cues and their relation to the task. This extended training might have allowed that the competition between the target and the task-irrelevant cues especially the ones from the same sensory modality is better resolved. In fact, in a subsequent study (Antono et al., 2022), we showed that after being exposed to the intra- and cross-modal reward cues that were predictive of the delivery of the reward upon correct performance, the visual discrimination was enhanced by previously rewarded cues of both modalities. This finding supports the idea that the duration of training and the history of reward delivery may influence the way that task-irrelevant previously rewarded stimuli affect the perceptual decisions (Jahfari and Theeuwes, 2017; Jahfari et al., 2020). Future studies will be needed to systematically investigate these factors.

In line with the behavioural results, we found that early visual areas within the anatomical boundaries of area V1 – V2 had a better representation of the tilt orientation of the target when the target was presented together with the high reward stimuli. Reward signals have been known to modulate the early sensory areas (*visual*: Bayer et al., 2017; Serences, 2008, *auditory*: Beitel, et al., 2003; Guo, et al., 2019, *somatosensory*: Pleger, et al., 2008). More specifically, it has been known that the early visual areas are sensitive to the reward magnitude (Serences, 2008; Weil et al., 2010; Arsenault et al., 2013) and timing (Shuler & Bear, 2006; Chubykin, et al., 2013). Importantly, the reward-driven modulations in our study were spatially specific and overlapped with the regions within the area V1-V2 that represented the visual target, in line with previous observations (Serences, 2008; Arsenault et al., 2013). In contrast, other studies have provided evidence that reward effects may rely on a combination of stimulus-specific and unspecific modulations, suggesting that reward learning in the visual system may be gated by mechanisms that are distinct from sensory processing (FitzGerald et al., 2013; Schiffer et al., 2014; Poort et al., 2015). Since in our design we did not manipulate the spatial location of stimuli and the delivery of rewards were halted during the test phase, we cannot infer the extent to which the spatial profile of reward-driven effects in our study reflects a general principle as opposed to a particular pattern imposed by our task design. Unravelling the spatial characteristics of reward-driven modulations from different sensory modalities is an important direction for future studies.

What mechanisms underlie the reward-driven enhancement of target representations in the early visual areas? We sought the answer to this question by first mapping the areas where the reward value was represented and thereafter testing different models of how reward information could be communicated between the valuation and early visual areas. Using a multivariate pattern classification approach, the lateral orbitofrontal cortex (OFC) was identified as a region that reliably encoded stimulus value independent of the sensory features of the reward associated stimuli. Previous studies have shown that this area plays a key role in representing the magnitude of rewards, especially when there is uncertainty in the appropriate course of action to be taken such as when previously rewarded responses should be suppressed (Elliott et al., 2000; O’Doherty et al., 2001). Furthermore, IPS and STS were identified by the value decoders which were sensitive to the sensory features of the reward stimuli (i.e., modality and location). IPS has been consistently linked to the processing of the goal-directed information and voluntary orienting towards a spatial location (Corbetta et al., 2000; Corbetta and Shulman, 2002; Serences and Yantis, 2007). Specifically, the coordinates observed in our study is close to the anterior part of the IPS with dense neuroanatomical connectivity with the frontal areas (Greenberg et al., 2012), suggesting that the modulation of IPS may be driven by the top-down signals from the frontal valuation areas. The superior temporal areas such as STS have been classically shown to be involved in the integration of information across sensory modalities (Calvert et al., 2001; Werner and Noppeney, 2010). Moreover, the role of this area in the integration of information has been shown to go beyond the multisensory processing and also include a general role in linking the sensory attributes of stimuli to the cognitive factors such as attention (Shapiro and Hillstrom, 2002), reward (Lim et al., 2013; Pooresmaeili et al., 2014) and affective and social processing (Beauchamp, 2015). Importantly, STS and IPS have been shown to have structural connectivity (Cavada and Goldman-Rakic, 1989) and form a network for attentional (Shapiro and Hillstrom, 2002) and multisensory processing (Werner and Noppeney, 2010), and additionally STS has been shown to communicate the reward-related information to the frontal valuation areas (Lim et al., 2013). Given these findings from the previous studies, the valuation areas identified by our approach constituted a plausible network, shown in **Figure 4**, to represent and communicate the information related to the reward value across the brain.

We next used an effective connectivity analysis to explicitly test how such a putative communication occurs. We tested different mechanisms that either relied on a direct or a mediated communication between the valuation and the early visual areas. This analysis supported a model which assumed the mediation of reward effects through attention and/or higher sensory areas. The communication between the valuation- and attention-related areas are aligned with the notion of attentional gated reward processing (Roelfsema and Van Ooyen, 2005). In line with this model, we found that when there was a need to discriminate the sensory features of reward- and task-related stimuli, as was the case when reward cues were from the same modality, the feedforward communication between the attentional and the valuation network was enhanced relative to when reward-related stimuli were highly distinct from the visual target (i.e. for cross-modal cues). On the other hand, previous studies have also proposed rewards to be a teaching signal for attention (Chelazzi et al., 2013), as the magnitude of reward determines the way that attention should be allocated in space. In line with this proposal, we found a general pattern across the sensory modalities where higher areas sent inhibitory feedback signals to upstream attentional and higher-order sensory areas, potentially in order to suppress the excessive allocation of attention and other processing resources to the task-irrelevant cues. Together, our findings show the fine-tuned mechanisms that underlie the regulation of attention and reward processing across the sensory modalities.

The pattern of connectivity modulations at the lower levels of the network shown in **Figure 4C** revealed further dissociations between the intra- and cross-modal rewards. Specifically, the communication from the IPS back to the early visual areas demonstrated a distinct pattern across intra- and cross-modal conditions. Whereas reward-related information was communicated from IPS directly to the early visual areas and elicited feedback inhibition, cross-modal cues required a mediation through a sensory-dependent area in the superior temporal areas and modulated the early visual areas through excitatory interactions. This pattern is in line with the findings of a previous study (Vakhrushev et al., 2021) where a dissociation between the reward-driven effects of previously rewarded intra- and cross-modal cues was found. Putatively, the feedback inhibition in case of the intra-modal reward cues reflects the down-weighting of the value of the task-irrelevant features of an object (i.e. the colors), which share processing resources with the target. In fact, recent studies have shown that at the level of area V1, processing of orientation and color is more inter-related than previously thought (Garg et al., 2019). This means that by regulating the processing of high reward colors through feedback inhibition, the early visual areas could better dedicate resources to the representation of the stimulus orientation. In contrast, in the cross-modal condition, there is little necessity to suppress the reward cues as they elicit a relatively weaker competition with the target at the level of the early visual areas. In fact, through enhancing the allocation of attention (Eimer and Driver, 2001) or the integration of the auditory tones and visual stimuli (Driver and Noesselt, 2008; Petro et al., 2017), a boost in the processing of cross-modal reward cues could potentially enhance the overall salience of the visual target at the level of early visual areas.

Altogether, the commonalities and dissociations between intra- and cross-modal rewards observed in the effective connectivity results point to two general patterns. Firstly, both reward types engage attentional areas and lead to a predominantly inhibitory feedback connectivity between the valuation and attentional areas. Hence, the regulation of information processing at the level of higher cognition seems to be modality-independent. Secondly, at the lower levels of hierarchy where reward-related information is relayed to the early visual areas, more dissociations between the intra- and cross-modal rewards emerge: not only do the cross-modal rewards additionally engage a higher-order sensory area (STS) but also they elicit an overall enhanced communication to and from the early visual areas, whereas intra-modal rewards evoked an overall inhibition. We interpret the dissociations between the intra- and cross-modal reward effects as a consequence of the differences in the way that they interact with the processing of the target at the level of early visual areas. Future studies will be needed to test whether a systematic relationship exists between the degree of overlap in neural mechanisms of task-relevant and reward-related features of stimuli and the way that perceptual decisions are influenced by the rewards.

Previous theoretical and empirical work has suggested a tight interaction between reward and attention (Roelfsema and Van Ooyen, 2005; Stanisor et al., 2013). In this vein, it has been suggested that attention and reward reinforcement (Seitz and Watanabe, 2009) can work as heuristics which help the visual system to determine the sensory features that are relevant. Similarly, Padmala and Pessoa (2011) discussed that reward information enables a coupling between the attentional and valuation networks. Specifically, comparing the functional connectivity of rewarded and not-rewarded trials (Padmala and Pessoa, 2011; Kinnison et al., 2012) they found that whereas in rewarded trials attentional and valuation mechanisms worked as an integrated system, in not-rewarded trials they worked more independently from each other. Extending these findings, we showed that the coordination of attention and valuation may additionally occur for previously rewarded stimuli and engage higher-order sensory areas such as STS. An important direction for future studies will be to examine whether the mode of interaction between reward-, attention- and sensory-related areas holds under different contexts for instance different attentional loads and contingencies of rewards to performance (Antono et al., 2022). Our hypothesis is that the visual system will engage both attention and reward systems as resources to learn and change its plasticity. However, depending on the availability and the reliability of the resources, it can flexibly rely on one system rather than the other. Furthermore, future studies will be needed to delineate whether the involvement of long-range interactions to and from the sensory areas is a general feature of reward-driven modulation of perception or a specific finding in the setting that we tested. It is conceivable that when rewards are consistently paired with the task-relevant features, they may induce long-lasting changes at the level of early sensory areas that locally enhance the processing of reward-related stimuli, as predicted by computational models (Wilmes and Clopath, 2019). In these cases, a long-term prioritization of reward-related stimuli is advantageous for the system as they could consistently lead to a behavioural gain for the organism. Quantifying the exact relationship between rewards’ availability and reliability and the degree to which they promote long-term plasticity in the early sensory areas is an exciting direction for future studies.

## Acknowledgements

We thank Tabea Hildebrand, Jana Znaniewitz, and Sanna Peter for their help with the data collection. We also thank Dr. Carsten Schmidt-Samoa and Dr. Peter Dechent for their technical assistance, and Prof. Melanie Wilke and Dr. Roberto Goya-Maldonado for their input and suggestions. This work was supported by an ERC Starting Grant (no: 716846) to AP.

## Authors’ contributions

JEA and AP conceptualized the project designed the task. JEA conducted the experiments. JEA, SD, RA, and AP analyzed the data. JEA and AP interpreted the results and wrote the first draft of the manuscript. All authors revised the manuscript. AP acquired funding.

## Supplementary Information

### Conditioning phase

To ensure that participants had learned reward-cue associations, we asked a question during and after the experiment. Based on these questionnaires, all participants could correctly identify which cue properties were associated with high compared to low reward magnitudes/ or were aware of the cue-reward associations. We further tested whether during the conditioning phase, reward predicting cues modulated participants’ behavior. Participants exhibited near perfect accuracy in localizing both visual and auditory stimuli (mean performance 99.9% and 96%±1%, respectively), with a consistent superiority of vision (F(1,32) = 18.36, *p* < .001, η_p_^2^ = 0.365). Similarly, participants’ responses were significantly faster in visual compared to auditory trials (F(1,32) = 70.94, *p* < .001, η_p_^2^ = 0.689). This result is in line with the superior performance of vision compared to audition in localization tasks. However, we found no significant main effect of reward on either accuracies (F(1,32) = 0.93, *p* = 0.34, η_p_^2^ = 0.028) or response times (F(1,32) = 0.29, *p* = 0.60, η_p_^2^ = 0.009), neither did we find an interaction between reward and modality on accuracies (F(1,32) = 0.71, *p* = 0.405, η_p_^2^ = 0.022) or response times (F(1,32) = 0.20, *p* = 0.66, η_p_^2^ = 0.006). This result is likely due to the fact that the task was already done at a near perfect level and performance had already reached a ceiling.

**Supplementary Figure 1.**
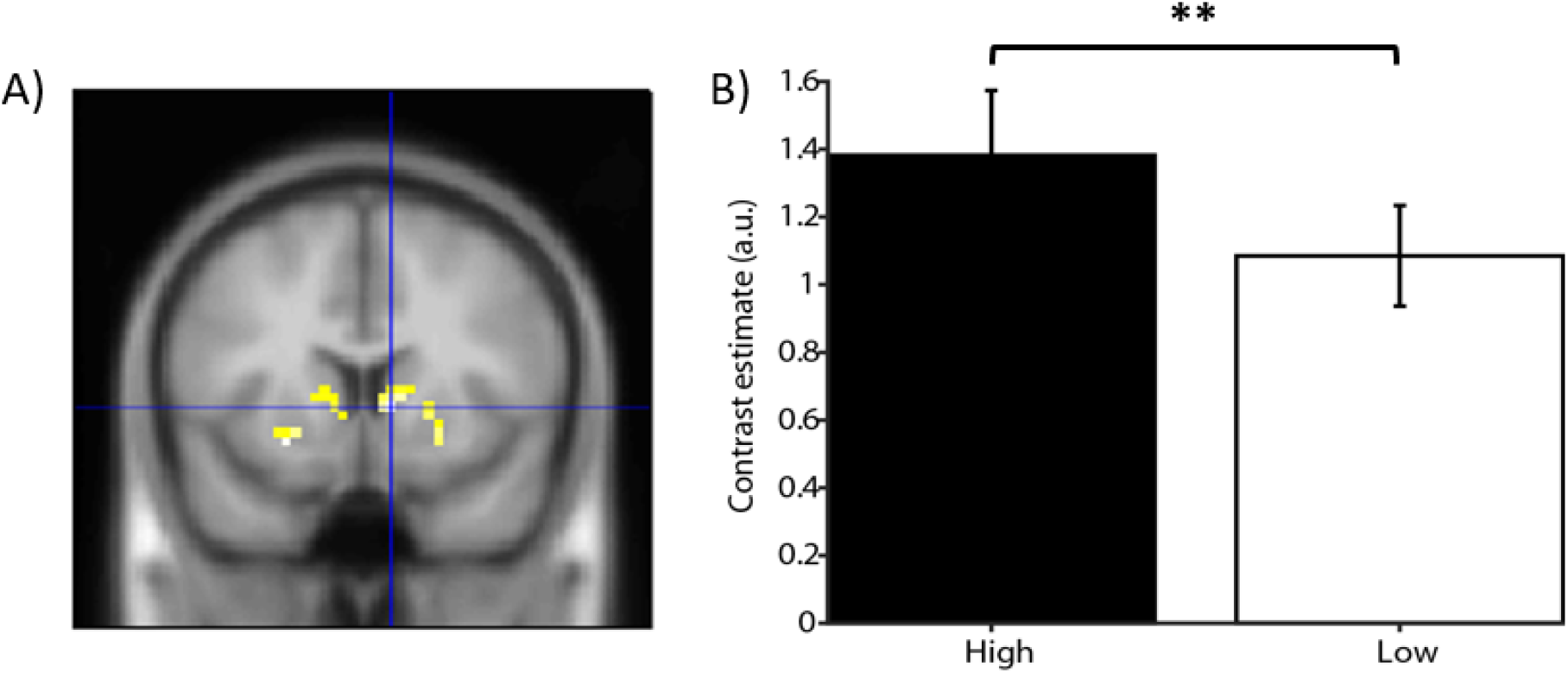
Main effect of reward (AH+VH>AL+VL, AH: Auditory High reward, VH: Visual High reward, AL: Auditory low reward and VL: Visual Low reward) during the conditioning phase. A) Contrast between high against low reward conditions, thresholded at *p* < .001 (uncorrected) with k = 10 and masked with an anatomical ROI encompassing the ventral striatum mask (i.e., Putamen, Caudate, and Globus Pallidus). Crosshair is at the peak activation xyz = [9 11 2]. B) Bar graphs depict the contrast estimates of high against low reward conditions. ** corresponds to *p* < 0.01 based on a paired sample *t*-test.

We next examined the reward-driven modulation of BOLD responses during the conditioning phase by inspecting the results of a mass-univariate analysis of fMRI data during this phase. As expected, several brain areas encompassing sensory (e.g. visual cortex) and reward-related areas (such as insula and the ventral striatum) demonstrated a strong modulation by rewards across modalities (i.e. for the contrast AH+VH > AL+VL, see **Supplementary Table S1**). These areas have been consistently shown to be involved in the processing of reward information during associative learning in previous studies (Schultz, 2000; Daniel and Pollmann, 2014), and indicate that participants did learn the association between cues and different reward magnitudes. For illustrative purposes, we examined the main effect of reward in a ROI encompassing the ventral striatum, an area that has been shown to be involved in learning of reward associations in a large body of previous literature (Schultz, 2000; Tremblay et al., 2009; Haber and Knutson, 2010). We found that higher reward magnitude modulated the ventral striatum compared to lower reward magnitude, as shown in **Supplementary Figure 1**. Moreover, we investigated further whether the reward effect had dependencies on the sensory modalities at this stage. To this end, we examined the interaction contrasts corresponding to a stronger reward effect in auditory (AH-AL > VH-AL) or visual (VH-VL > AH-AL) stimuli. We observed that a right lateralized Caudate [15 -22 26] cluster that was modulated stronger by auditory compared to visual cues, whereas no activations were found for the opposite contrast (VH-VL > AH-AL). This result demonstrated that auditory stimuli elicited stronger activations in reward-related areas such as Caudate, suggesting that the sensory modality of rewards could be to some extent dissociated in the reward-related areas. Moreover, this result may also be due to a higher saliency of auditory compared to visual stimuli, as the Caudate has been known to encode saliency of the sensory stimuli (Zink et al., 2006).

**Supplementary Table 1.**
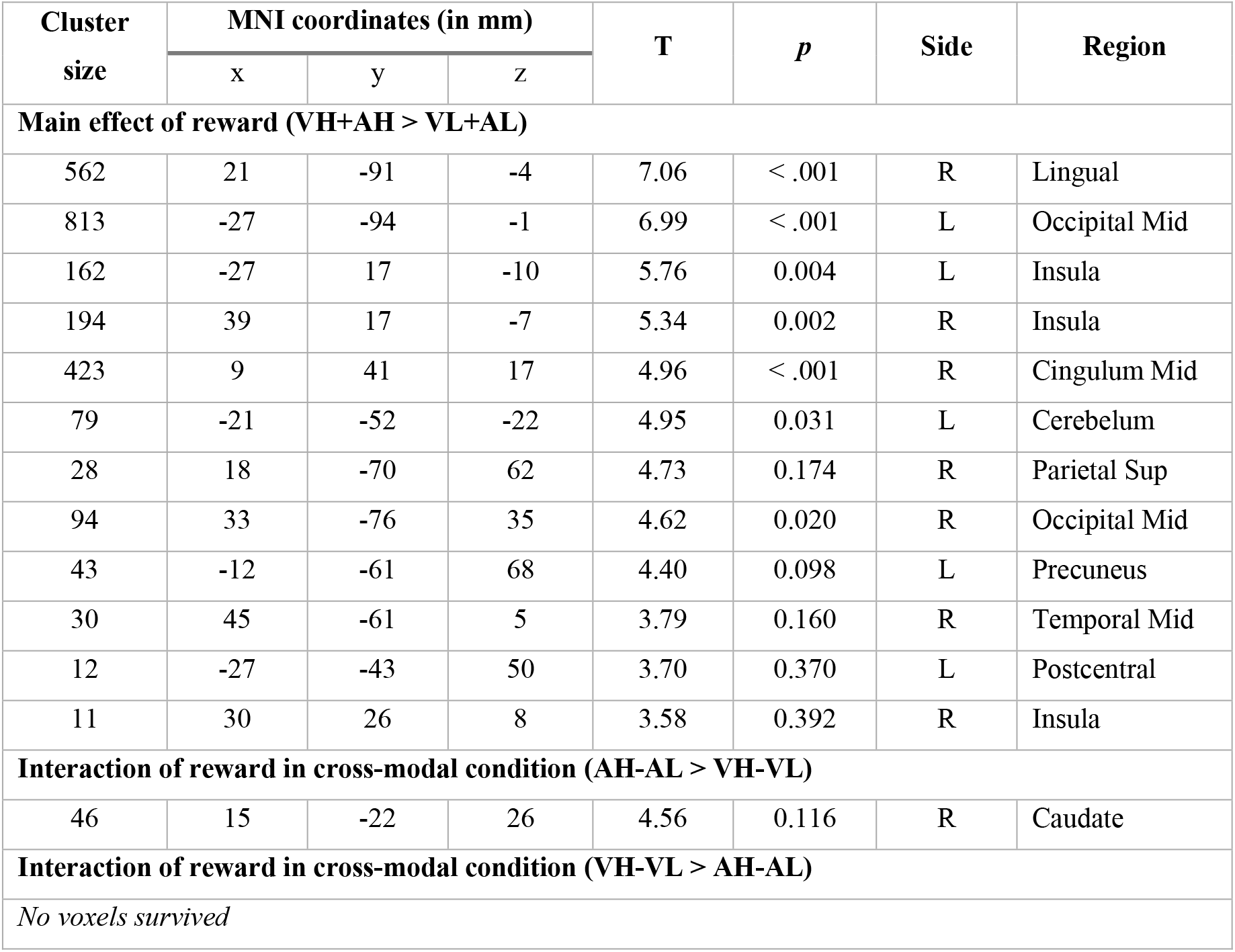
Whole-brain analysis result during conditioning phase with uncorrected threshold of p < .001 and extent threshold of k = 10.

| Cluster size | MNI coordinates (in mm) |  |  | T | *p* | Side | Region |
| --- | --- | --- | --- | --- | --- | --- | --- |
|  | x | y | z |  |  |  |  |
| Main effect of reward (VH+AH > VL+AL) |  |  |  |  |  |  |  |
| 562 | 21 | -91 | -4 | 7.06 | < .001 | R | Lingual |
| 813 | -27 | -94 | -1 | 6.99 | < .001 | L | Occipital Mid |
| 162 | -27 | 17 | -10 | 5.76 | 0.004 | L | Insula |
| 194 | 39 | 17 | -7 | 5.34 | 0.002 | R | Insula |
| 423 | 9 | 41 | 17 | 4.96 | < .001 | R | Cingulum Mid |
| 79 | -21 | -52 | -22 | 4.95 | 0.031 | L | Cerebellum |
| 28 | 18 | -70 | 62 | 4.73 | 0.174 | R | Parietal Sup |
| 94 | 33 | -76 | 35 | 4.62 | 0.020 | R | Occipital Mid |
| 43 | -12 | -61 | 68 | 4.40 | 0.098 | L | Precuneus |
| 30 | 45 | -61 | 5 | 3.79 | 0.160 | R | Temporal Mid |
| 12 | -27 | -43 | 50 | 3.70 | 0.370 | L | Postcentral |
| 11 | 30 | 26 | 8 | 3.58 | 0.392 | R | Insula |
| Interaction of reward in cross-modal condition (AH-AL > VH-VL) |  |  |  |  |  |  |  |
| 46 | 15 | -22 | 26 | 4.56 | 0.116 | R | Caudate |
| Interaction of reward in cross-modal condition (VH-VL > AH-AL) |  |  |  |  |  |  |  |
| No voxels survived |  |  |  |  |  |  |  |

### Reward modulation on the early visual areas overlapped with target processing areas

To find a region in the early visual areas that was specifically responsive to the visual target (i.e., was target-specific), we inspected the univariate contrast of Neutral cues > Baseline (corrected at pFWE < 0.05, k = 0, cyan region in **Supplementary Figure 2**). The effect of rewards on the representation of visual target shown in **Figure 2B** of the main text, spatially overlapped with the target-specific regions identified by the above contrast (magenta activations in **Supplementary Figure 2)**. Specifically, reward-driven modulations (right n of voxels = 19; left n of voxels = 14) were small-volume corrected within a mask comprising of target-specific activations (right n of voxels = 14; left n of voxels = 13, *p* uncorrected < 0.005, k = 10, see **figure S2**), indicating that most of the activated voxels correspond to the target-specific regions. Nevertheless, we took the whole cluster within V1-V2 anatomical mask for the effective connectivity analysis.

**Supplementary Figure 2.**
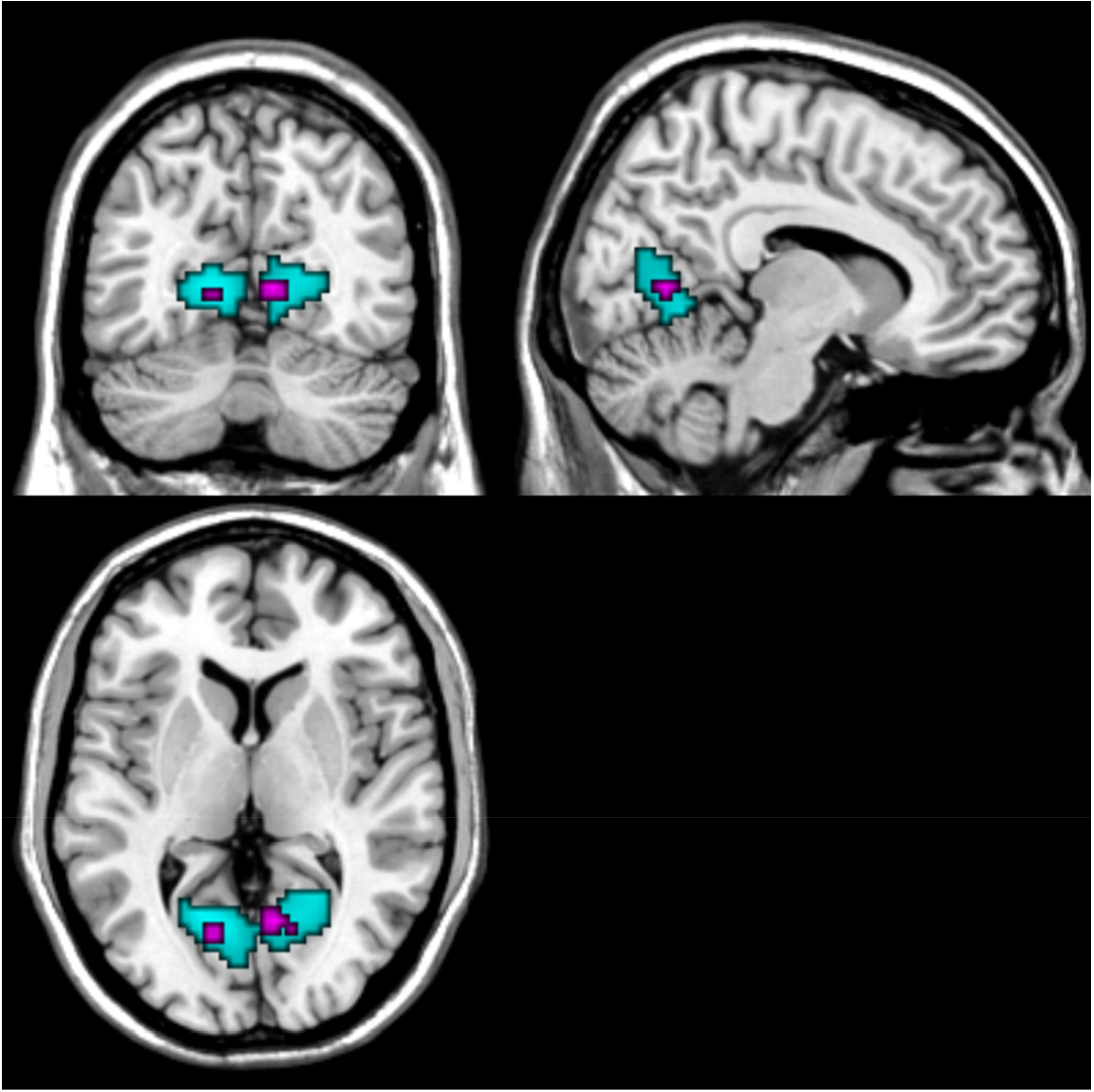
Reward facilitation in the early visual areas (masked with V1-V2 anatomical mask) overlapped with areas responsible for processing the target cue. Cyan color shows the response magnitude of target processing (Neutral vs Baseline) thresholded at *p*FWE < .05 and k = 0. Magenta color shows the rewardd-driven facilitation effect in visual areas thresholded at uncorrected *p* < .005, k = 10. The cursor is at xyz=[9 −64 5].

### Whole-brain results of the *value decoders*

**Supplementary Figure 3.**
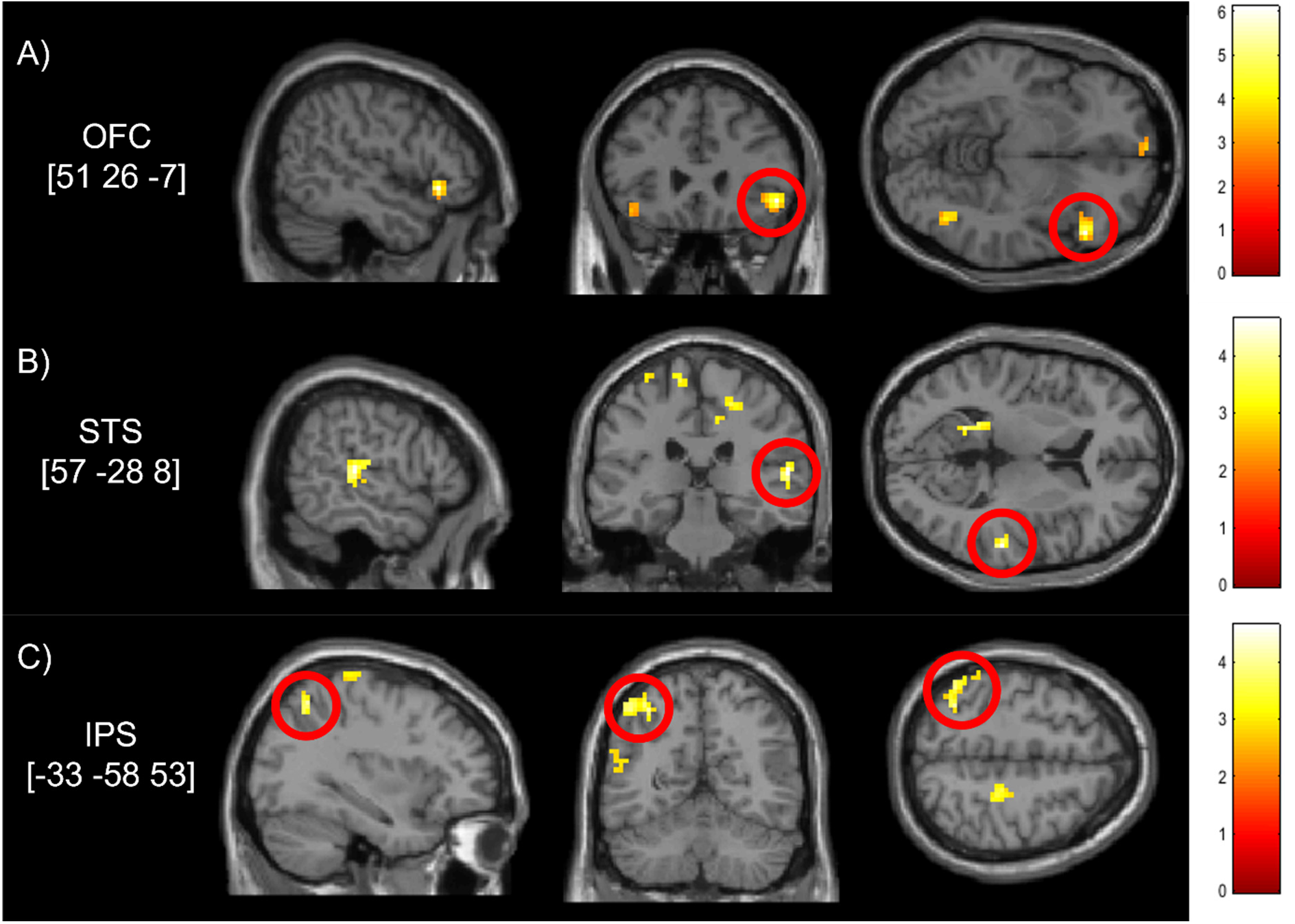
Whole-brain results of the value-decoders depicting sagittal, coronal, and the axial view for: A) lateral orbitofrontal areas xyz = [51 26 −7] in the right hemisphere from *value decoder 1*. B) The left anterior intraparietal areas xyz = [−33 −58 53] and C) The right superior temporal areas xyz = [57 −28 8] detected by the *value decoder 2* across sensory modalities. These ROIs were taken further to the effective connectivity analysis. All images were thresholded at uncorrected p < .005, k = 20. The cursor is at the peak activities of each corresponding ROI coordinates written in brackets.

Regions of interests (ROI) extracted from the whole-brain results of the *value decoder*_*1*_ and *value decoder*_*2*_ (see Materials and Methods), where the right lateral orbitofrontal areas had the strongest modulation in the *value decoder1* (see **figure S3**). Other areas such as the Caudate, Cerebelum, and the left lateral orbitofrontal areas were also modulated by reward (see **Table 1**). Moreover, *value decoder2* showed the right superior temporal areas had the strongest reward modulation. Interestingly, areas that have been linked to attentional processing in the anterior intraparietal areas (Corbetta and Shulman, 2002) also demonstrated as the largest areas modulated by reward. The time-series of these areas were extracted for effective connectivity between the reward-related areas in the lateral orbitofrontal cortex and the early visual areas.

